# lncRNA-RMST functioned as a SOX2 transcription co-regulator to regulate miR-1251 in the progression of Hirschsprung’s disease

**DOI:** 10.1101/2020.09.23.309930

**Authors:** Lingling Zhou, Zhengke Zhi, Pingfa Chen, Zhonghong Wei, Chunxia Du, Binyu Wang, Xiang Fang, Weibing Tang, Hongxing Li

**Affiliations:** Department of Pediatric Surgery, Children’s Hospital of Nanjing Medical University, Nanjing, 210008, China; Soochow University Affiliated Children’s Hospital, Suzhou, 215128, China; Yancheng city NO.1 People’s hospital, Yancheng, 224006, China

**Author notes:** These authors have contributed equally to this work (Lingling Zhou, Zhengke Zhi, Pingfa Chen). Correspondence to: Hongxing Li.

**Keywords:** Hirschsprung’s disease, lncRNA-RMST, miR-1251, SOX2, AHNAK

## Abstract

Hirschsprung’s disease (HSCR) is a congenital disorder characterized by the absence of enteric neural crest cells (ENCCs). Non-coding RNAs including long non-coding RNAs (lncRNAs) and microRNAs (miRNAs) have been authenticated as important regulators of biological functions. We conducted a microarray analysis and found lncRNA Rhabdomyosarcoma 2-associated transcript (RMST) was down-regulated in the stenotic segment of HSCR patients. MiR-1251 is transcribed from the intron region of RMST and was also low-expressed. When the expression of RMST or miR-1251 was reduced, the cell proliferation and migration were attenuated. However, RMST didn’t affect the expression of miR-1251 directly found in this study. Through bioinformatic analysis, transcription factor SOX2 was predicted to bind to the promoter region of miR-1251 which was confirmed by CHIP assay. Herein, we demonstrated that RMST exerted as a co-regulator of SOX2 to regulate the expression of miR-1251. Furtherly, AHNAK was proved to be the target gene of miR-1251 in this study. Taken together, we revealed the role of RMST/SOX2/miR-1251/AHNAK pathway in the occurrence of Hirschsprung’s disease and provided a potential therapeutic target for this disease.

**SUMMARY STATEMENT:** Hirschsprung disease (HSCR) is characterized by a deficit in enteric neurons, however, the underlying mechanism remains unclear. This study revealed the role of lnc-RMST during the occurrence of HSCR.

## 1. INTRODUCTION

Hirschsprung disease (HSCR), a common enteric neuropathy, is characterized by the absence of gangliocytes in the distal colon(Jaroy et al., 2019; Sergi, 2015). During 5 to 12 weeks of embryogenesis, enteric neural crest cells (ENCCs) failed to migrate and proliferate might cause this disease (Bergeron et al., 2013). HSCR usually attacks about 1/5000 neonates, while the incidence rate of females is about a quarter of males. (Wester and Granstrom, 2017). Current etiological studies show that HSCR is a complicated disorder involving multiples genetic factors(McKeown et al., 2013). Genes including RET, GDNF, GFRA1, EDNRB and PHOX2B have been confirmed to be involved in HSCR (Tam, 2016; Zhao et al., 2019). However, these genes could only explain partly, so further research is needed.

With longer than 200 nucleotides, long noncoding RNAs (lncRNAs) are increasingly considered to be the main players in governing basic biological processes by affecting gene expression at nearly all levels.(Shen et al., 2019; Xu et al., 2020). As reported before, various lncRNAs can regulate cell proliferation and migration. For instance, lncRNA TPTEP1 could inhibit the non-small cell lung cancer (NSCLC) cells to proliferate through abating miR-328-5p expression(Cao et al., 2020). In addition, in renal cell carcinoma, lncRNA00312 attenuated cell proliferation and migration obviously(Zeng et al., 2020). However, the study about lncRNA functioned in HSCR is rarely reported.

In order to explore the role of lncRNA in the occurrence of HSCR, a microarray was conducted in this study, and we found lncRNA Rhabdomyosarcoma 2-Associated Transcript (RMST) was significantly low expressed in the aganglionic bowels compared with the normal ones. RMST has been discovered to be essential in neuronal differentiation(Cheng et al., 2020; Tang et al., 2015). According to a report, RMST promoted activation of microglial cells by activating TAK1-mediated NF-κB signaling(Sun et al., 2019). Considering the effects of RMST on nervous system and the low-expressed of RMST in HSCR, we aimed to reveal its roles during the procedure of HSCR. Furtherly, miR-1251 was transcribed from the same genomic site as RMST and was also low-expressed in HSCR diseased bowel. However, we found RMST didn’t regulate its intron gene miR-1251 independently in this study. There may be other regulatory mechanisms to be explored.

Sex determining region Y (SRY)-box 2 (SOX2) functions a transcription factor is implicated in transcriptional regulation(Collignon et al., 1996; Schepers et al., 2002). It has also been discovered to regulate miRNAs expression(Liu et al., 2017). Interestingly, SOX2 is closely related to the nervous system, such as the terminal differentiation of postmitotic olfactory neurons was regulated by SOX2 directly (Alqadah et al., 2015). Through bioinformatics analysis, it was found SOX2 might bind to the promoter of miR-1251. Numerous evidences have also indicated that the down-regulation of SOX2 attenuated cell growth and migration(Sannino et al., 2019; Schaefer and Lengerke, 2020). Thus, SOX2 probably be related to the development of neural crest cells during HSCR by regulating the expression of miR-1251. Furthermore, according to previous report, RMST could interact with SOX2 and then enhance its regulation on downstream genes(Ng et al., 2013). Therefore, we hypothesized that RMST might function as a SOX2 transcription co-regulator to regulate the downstream miR-1251 and participate in the proceeding of HSCR.

## 2. MATERIALS AND METHODS

### 2.1 Clinical information

This study was approved by the Institutional Ethics Committee of Nanjing Medical University (NJMU Birth Cohort), and the experiments were carried out according to approved guidelines. 32 stenotic colon tissues were collected from patients accepted radical operation of HSCR in Children’s Hospital of Nanjing Medical University from January 2011 to August 2014. 32 controls matched with cases on age and gender were randomly picked out from isolated patients on account of intussusception or incarcerated and strangulated inguinal hernia without the ischemia or necrosis parts. Tissues were harvested and stored at −80 °C immediately after surgery. All HSCR patients were diagnosed through pathological analysis. Finally, written informed consent from all participants were obtained.

### 2.2 Quantitative Real-time Polymerase Chain Reaction (qRT-PCR)

To isolate total RNA from tissues and cells, Trizol reagent (Invitrogen Life Technologies Co, USA) was applied. QRT-PCR was employed to detect RMST, miR-1251 and AHNAK expression. TaqMan^®^ MicroRNA Assays (Applied Biosystems, USA) was used to test miR-1251 expression. GAPDH and U6 was applied as an internal control for mRNA and miRNA detection, respectively. Roche LightCycler480 (Roche, Switzerland) was used to perform qRT-PCR depending on the manufacturer’s protocol. Primer sequences were showed in Table 1.

**Table 1.**
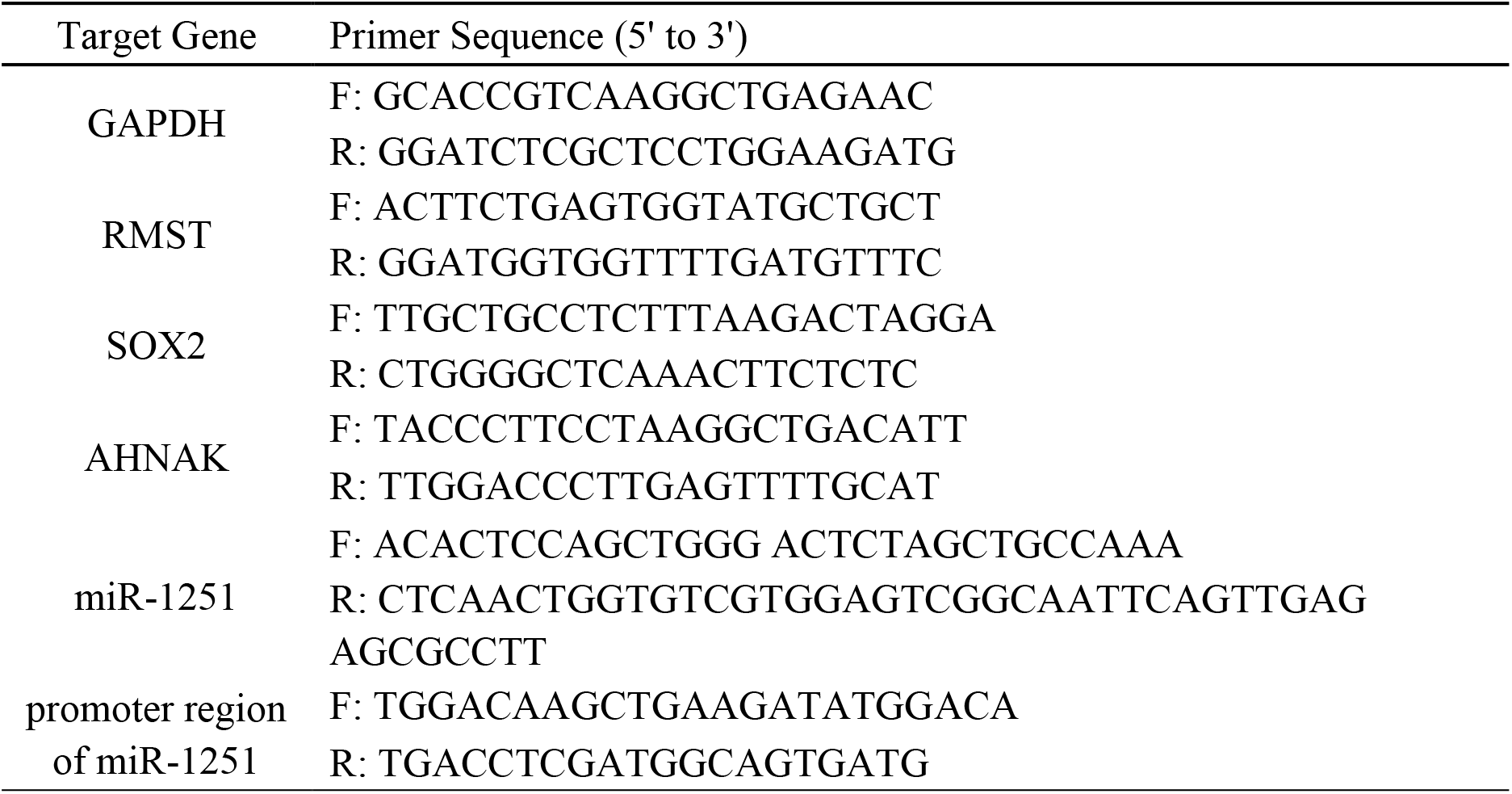
Primer Sequences for Quantitative RT-PCR

### 2.3 Western Blotting

RIPA lysis buffer (Beyotime, Shanghai, China) was applied to extract total proteins from colon tissues and cultured cells. BCA Protein Assay Kit (Beyotime, Shanghai, China) was used to detect protein concentration. The same amount of total proteins was isolated in 10% SDS-PAGE, transferred to PVDF membranes and then sealed for 1 h. At 4°C, primary antibodies were used for incubation overnight. Afterwards, corresponding secondary antibodies were added for 2 h at 25°C. Finally, the membranes were exposed via ECL and Western blot detection reagents (Thermo Fisher Scientific, MA, USA). Antibodies including anti-AHNAK (SC134252), anti-SOX2 (SC17320X) and anti-GAPDH (SC47724) were obtained from Santa Cruz (CA, USA). The corresponding secondary antibodies were obtained from Beyotime (Shanghai, China).

### 2.4 Chromatin Immunoprecipitation (ChIP)

By using ChIP Assay Kit (Thermo Fisher Scientific, Shanghai, China), ChIP was implemented in accordance with the operating instructions. Firstly, cross-linked chromatin was sonicated into around 200 bp to 1000 bp fragments. Anti-SOX2 was used to immunoprecipitate the chromatin. Goat immunoglobulin G (IgG, ab172730, Abcam, USA) was applied to be the negative control. PCR was performed using SYBR Green Mix (Takara Bio, Japan). The primer sequences were shown in Table 1.

### 2.5 Cell culture and transfection

SH-SY5Y and 293T cell lines were acquired from ATCC. Cells were cultured at 37°C, 5% CO_2_ condition using DMEM (Hyclone, USA) culture medium containing 10% FBS, 100 U/mL penicillin, and 100 μg/mL streptomycin. The inhibitor of miR-1251, siRNA of RMST, SOX2 and the corresponding negative controls were synthesized by Genechem (Shanghai, China). Transfection experiments were conducted by using Lipofectamine 2000 Reagent (Invitrogen Life Technologies Co, USA).

### 2.6 Cell proliferation assay

To test the cell viability, cell counting kit-8 (CCK-8, Dojindo, Japan) was employed. After transfection, cells were cultured in 96-well plates for 24-48 h and then cells were incubated with CCK-8 reagent the for 1-2 h. Eventually, the OD value at 450 nm was detected by the TECAN infinite M200 Multimode microplate reader (Tecan, Mechelen, Belgium). Each assay was conducted independently in triplicate.

### 2.7 Cell migration assay

Transwell chambers were placed above a 24-well plate. After transfection around 24-48h, cells were resuspended with serum-free medium to 1×10^6^ cells/ml. About 100μl cell suspension was seeded to the upper chamber. 500μL of complete culture medium containing FBS was added to the lower chamber. 24-48h later, 4% paraformaldehyde was applied to fix the lower chamber cells and then crystal violet staining solution was used to stain cells. Cells migrated to the lower chamber were counted and imaged using an inverted microscope (×20). All experiments were conducted in triplicate.

### 2.8 Dual-luciferase reporter assay

The predicted 3’-UTR sequence of AHNAK binding to miR-1251 and the mutated sequence were inserted into the pGL3 promoter vector (Genechem, Shanghai, China) named pGL3-AHNAK-WT and pGL3-AHNAK-MUT. For reporter assay, cells were planted into 24-well plates and transfected with 100ng of pGL3-AHNAK-WT and pGL3-AHNAK-MUT, 50nM miR-1251 mimics and negative control using Lipofectamine 2000. Renilla luciferase vector pRL-SV40 (5 ng) was transfected into cells as control. Based on the obtained ratio, the activation degree of target reporter genes in different sample was compared.

### 2.9 Statistical analysis

GraphPad Prism 7.0 (GraphPad Software, USA) was adopted to analyze data. Between two groups, *t*-test was applied to determine the statistically significant differences, while the comparison among multiple groups was performed via one-way ANOVA. All data were presented as the mean± SEM. P < 0.05 was considered as statistically significant.

## 3. RESULTS

### 3.1 Down-regulation of RMST and miR-1251 in HSCR patients

To verify the expression levels of RMST, qRT-PCR was employed. RMST was markedly down-regulated in ganglia-free intestinal segment compared with normal controls as result showed (Fig. 1A). In order to detect whether RMST could affect cell migration and proliferation, Transwell and CCK8 assays were conducted. As the results showed, after transfected with RMST siRNA, the migrated and proliferated cells were obviously fewer than the normal control in both cell lines. (Fig. 1B). We also discovered the RMST intronic miR-1251 was down-regulated in aganglionic tracts (Fig. 1C). When cells transfected with miR-1251 inhibitor, the cell migration and proliferation was attenuated (Fig. 1D).

**Fig. 1.**
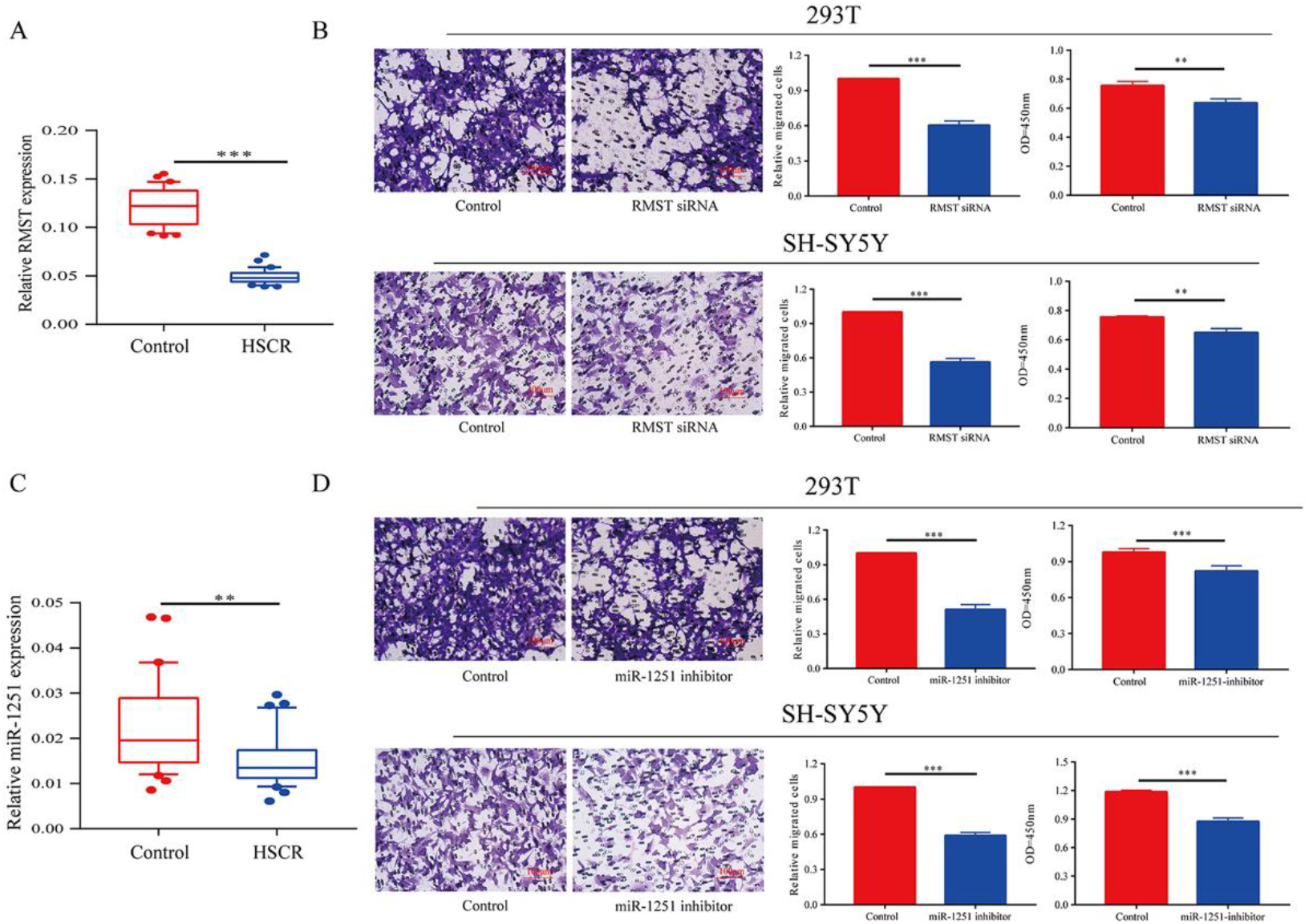
Down-regulation of RMST and miR-1251 in HSCR patients. **A** The expression of RMST was down-regulated obviously in stenosis tracts than control ones. **B** Transwell and CCK-8 assays showed cell migration and proliferation were attenuated when RMST was down-regulated. **C** The expression of miR-1251 was down-regulated obviously in stenosis tracts than control ones. **D** Cell proliferation and migration were inhibited obviously when transfected with miR-1251 inhibitor. \*\**P*<0.01, \*\*\**P*<0.001

### 3.2 miR-1251 was transcriptionally regulated by SOX2

Because miR-1251 is transcribed from the same genomic locus as RMST, we suspected that RMST might influence the expression level of miR-1251. To verify it, we knocked down RMST in SH-SH-SY5Y and 293T cells and then measured miR-1251 levels by qRT-PCR, however, there was no significant changes on miR1251 expression level in both cell lines, indicating that RMST was not a precursor transcript for it (Fig. S1 A). Furtherly, we employed bioinformatics approach Promoter Scan to predict the transcription promoter of miR-1251. SOX2 was predicted to bind with the 2kbp upstream promoter region of miR-1251 (Fig. S1 F). To confirm the combinative relationship between SOX2 and miR-1251, ChIP experiment was performed in the 293T cells. The result confirmed that SOX2 could bind to the promoter region of miR-1251 (Fig. 2A). Furtherly, when we abated the level of SOX2, miR-1251 was down-regulated obviously (Fig. 2B). Additionally, SOX2 was found down-regulated at mRNA and protein levels in HSCR patients than normal controls. (Fig. 2C, D). Based on this, we supposed that SOX2 influenced cell migration and proliferation by regulating miR-1251. As expected, when cells were transfected with SOX2 siRNA, cell proliferation and migration was attenuated, while upregulating miR-1251 could reverse it partly (Fig. 2E-G).

**Fig. 2.**
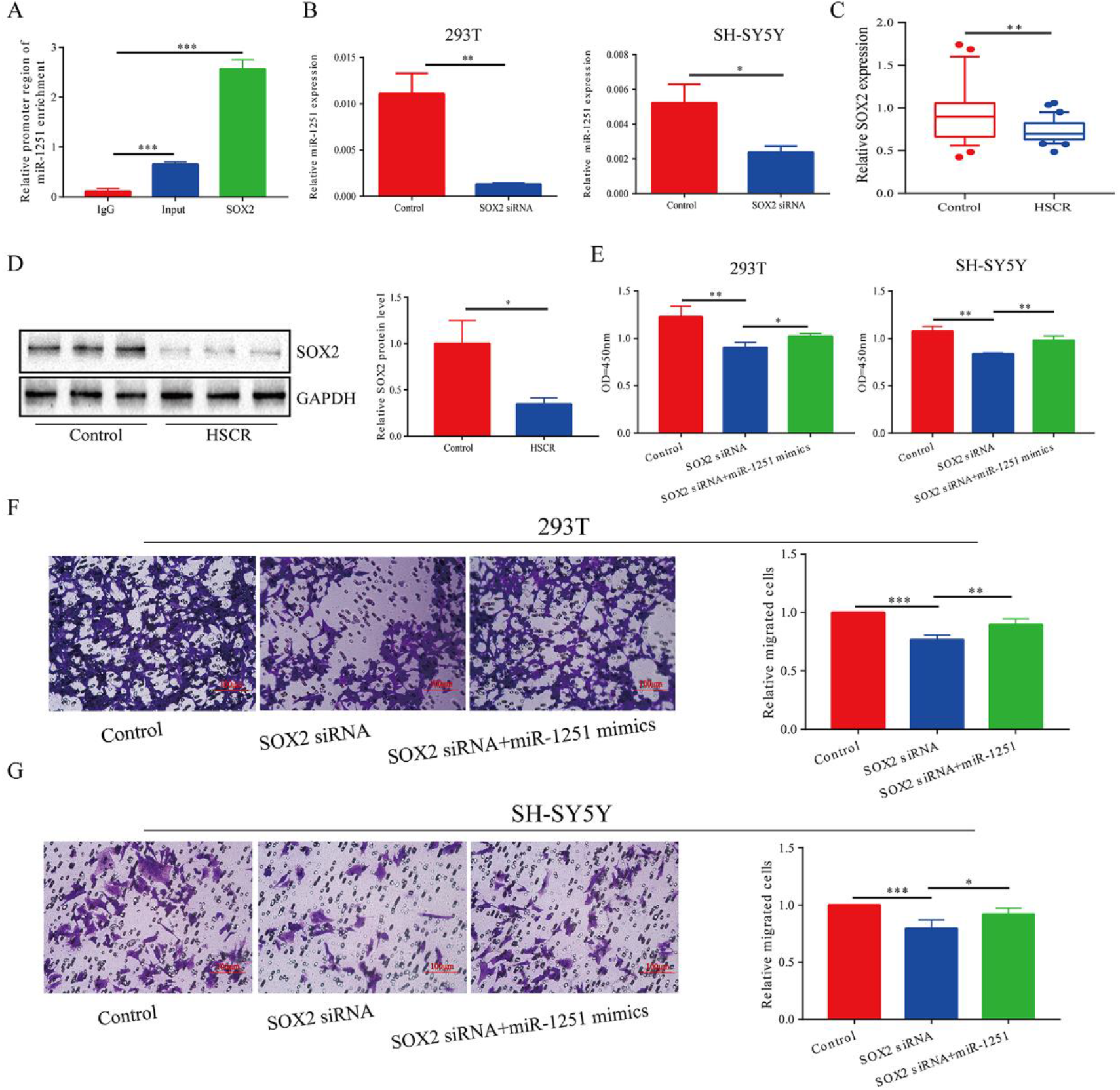
miR-1251 was transcriptionally regulated by SOX2. **A** CHIP assay showed SOX2 could bind to the promoter region of miR-1251. **B** After down regulating the expression of SOX2, the expression of miR-1251 was lower than control group. **C and D** The mRNA and protein expression of SOX2 in HSCR patients. **E-G** When transfected with si-SOX2, cell migration and proliferation were inhibited and the up-regulation of miR-1251 could partly reversed it. \**P*<0.05, \*\**P*<0.01, \*\*\**P*<0.001

### 3.3 RMST functioned as a co-regulator of SOX2

As reported before, RMST could also combine with SOX2 and then enhanced the transcriptional function of SOX2. In this study, we demonstrated that RMST could bind with SOX2 using RIP assay (Fig. 3A). Therefore, we forecasted that miR-1251 was transcriptionally regulated by SOX2 and RMST could strengthen this effect. When we knocked down the expression of SOX2 and both SOX2 and RMST in 293T cell, respectively, the expression of miR-1251 was detected. As expected, miR-1251 was down-regulated after cells transfected with SOX2 siRNA and was much lower in cells co-transfected with RMST siRNA and SOX2 siRNA (Fig. 3B). So, whether RMST exerted its roles through SOX2/miR-1251 axis? Through CCK-8 and Tranwell assays, we found that when the expression of RMST and SOX2 were both knocked down, the cell proliferation and migration were more weakened than just down-regulated RMST or SOX2 alone. Meanwhile, the upregulation of miR-1251 partly reversed the co-function of si-RMST and si-SOX2 (Fig. 3C, D). These results revealed that RMST might function via acting as a co-regulator of SOX2 to regulate downstream gene miR-1251.

**Fig. 3.**
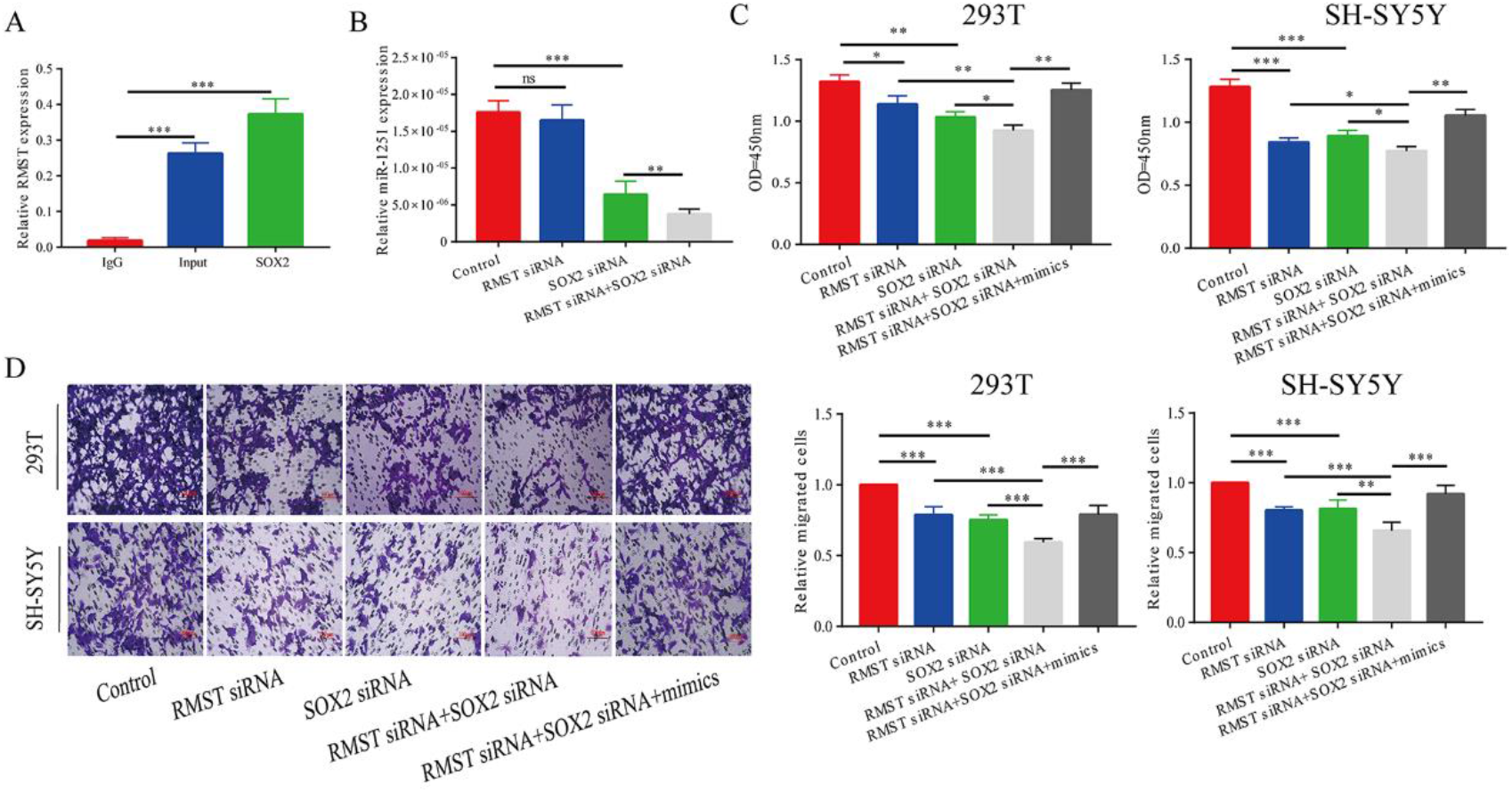
RMST functioned as a co-regulator of SOX2. **A** RIP assay showed RMST could bind to SOX2. **B** RMST functioned as a co-regulator of SOX2 and promoted the regulation effect of SOX2 on miR-1251. **C and D** CCK-8 and Tranwell assays revealed that when the expression of RMST and SOX2 were both knockdown, the cell proliferation and migration were more weakened than just down-regulated RMST or SOX2 alone. \**P*<0.05, \*\**P*<0.01, \*\*\**P*<0.001

### 3.4 AHNAK was the target gene of miR-1251

To predict the downstream target gene of miR-1251, bioinformatical analysis was employed and we found miR-1251 might bind to the 3’UTR of AHNAK (Fig. 4A). As dul-luciferase reporter assay showed, compared with the control group, the luciferase activity was significantly decreased when cells co-transfected with miR-1251 mimics and pGL3-AHNAK-WT plasmids demonstrating the relationship between miR-1251 and AHNAK (Fig. 4B). Furtherly, miR-1251 inhibitor was transfected in 293T cells. After 24 hours, AHNAK mRNA and protein levels were found up-regulated in 293T cells (Fig. 4C, D). By qRT-PCR, AHNAK was demonstrated up-regulated remarkably in aganglionic tract compared with normal controls (Fig. 4E). The protein level of AHNAK was detected furtherly, and was fit with its mRNA expression level (Fig. 4F). Meanwhile, we explored whether miR-1251 regulated cell function through AHNAK. As rescue experiment results showed, the reduction of AHNAK could partly reverse the influence of miR-1251 inhibitor on both cell migration and proliferation (Fig. 4G-I).

**Fig. 4.**
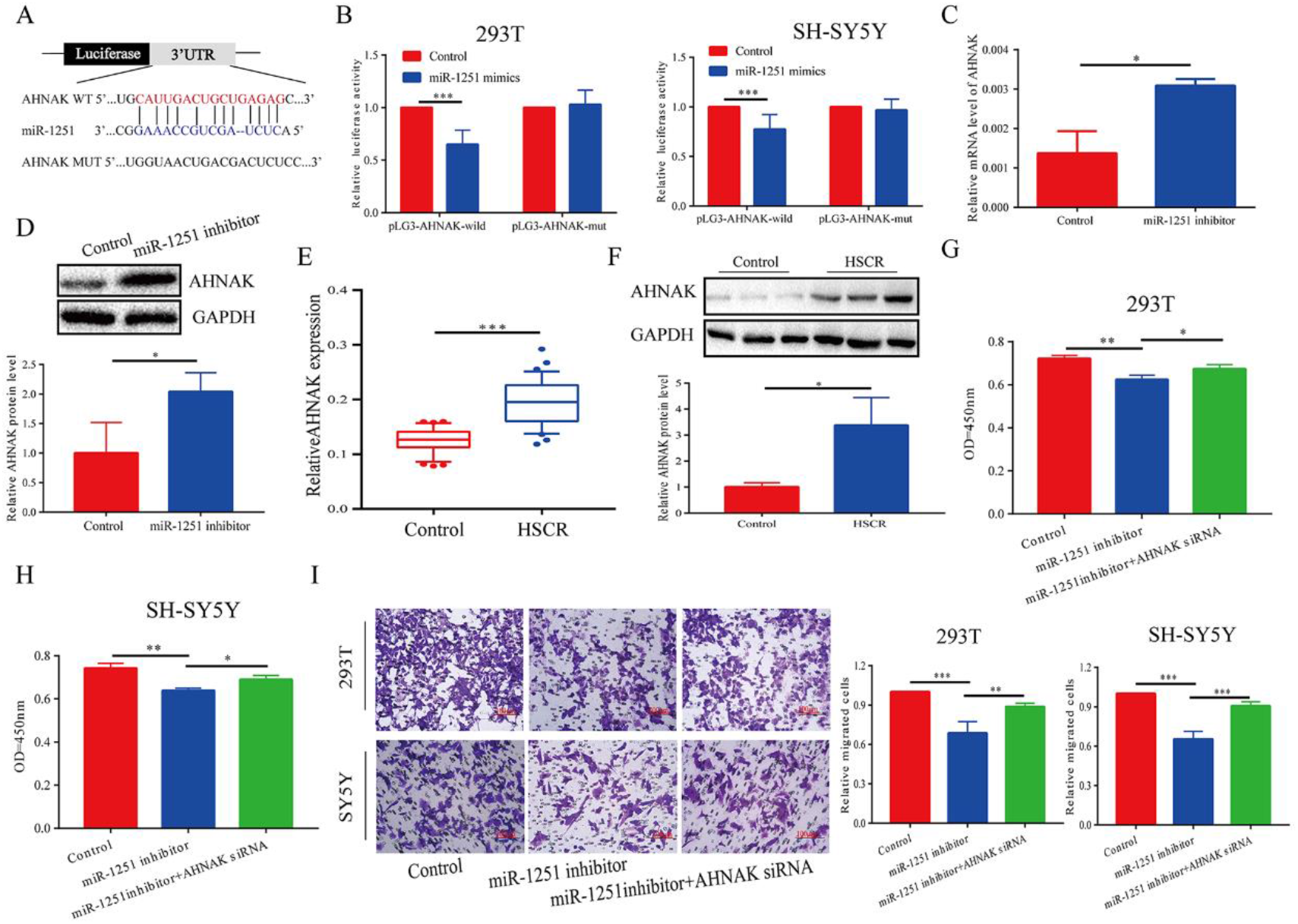
AHNAK was the target gene of miR-1251. **A** miR-1251 and AHNAK has potential binding sites. **B** Dul-luciferase reporter gene assay confirmed the binding relationship between miR-1251 and AHNAK in 293T and SY5Y cells. **C and D** When miR-1251 was knocked down, the mRNA and protein level of AHNAK was up-regulated. **E and F** The AHNAK mRNA and protein expression level in stenosis tracts was higher than control tracts. **G-I** The down-regulation of AHNAK could partly reverse the influence of miR-1251 inhibitor on cell migration and proliferation. \**P*<0.05, \*\**P*<0.01, \*\*\**P*<0.001

### 3.5 RMST played as a SOX2 transcription co-regulator to inhibit miR-1251 and raise AHNAK expression

Combined with above results, we presumed that RMST enhanced the regulation of SOX2 to miR-1251 and then promoted the expression of AHNAK, finally, affecting the proliferation and migration of neural cells. Firstly, the mRNA and protein expression of AHNAK was measured via qRT-PCR and western blot in every group respectively. There was no significant difference found in AHNAK expression between the RMST low expression group and the control group, however, the mRNA and protein expression of AHNAK was increased after SOX2 was down regulated. Furthermore, the expression of AHNAK was much lower in “RMST siRNA+SOX2 siRNA” group than in “SOX2 siRNA” group. which confirmed that RMST, as a SOX2 transcription co regulatory factor, upregulated the expression of AHNAK (Fig. 5A, B). In order to further confirmed the mechanism of its function through AHNAK, RMST low expression group, SOX2 low expression group, RMST low expression + SOX2 low expression group, RMST low expression + SOX2 low expression + AHNAK low expression group and control group were set up. CCK-8 and Transwell assays were applied to detect cell proliferation and migration abilities of every group. The results showed that the combined inhibition of both RMST and SOX2 low expression on cell proliferation and migration could be partially alleviated by simultaneously downregulating the expression of AHNAK (Fig. 5C, D).

**Fig. 5.**
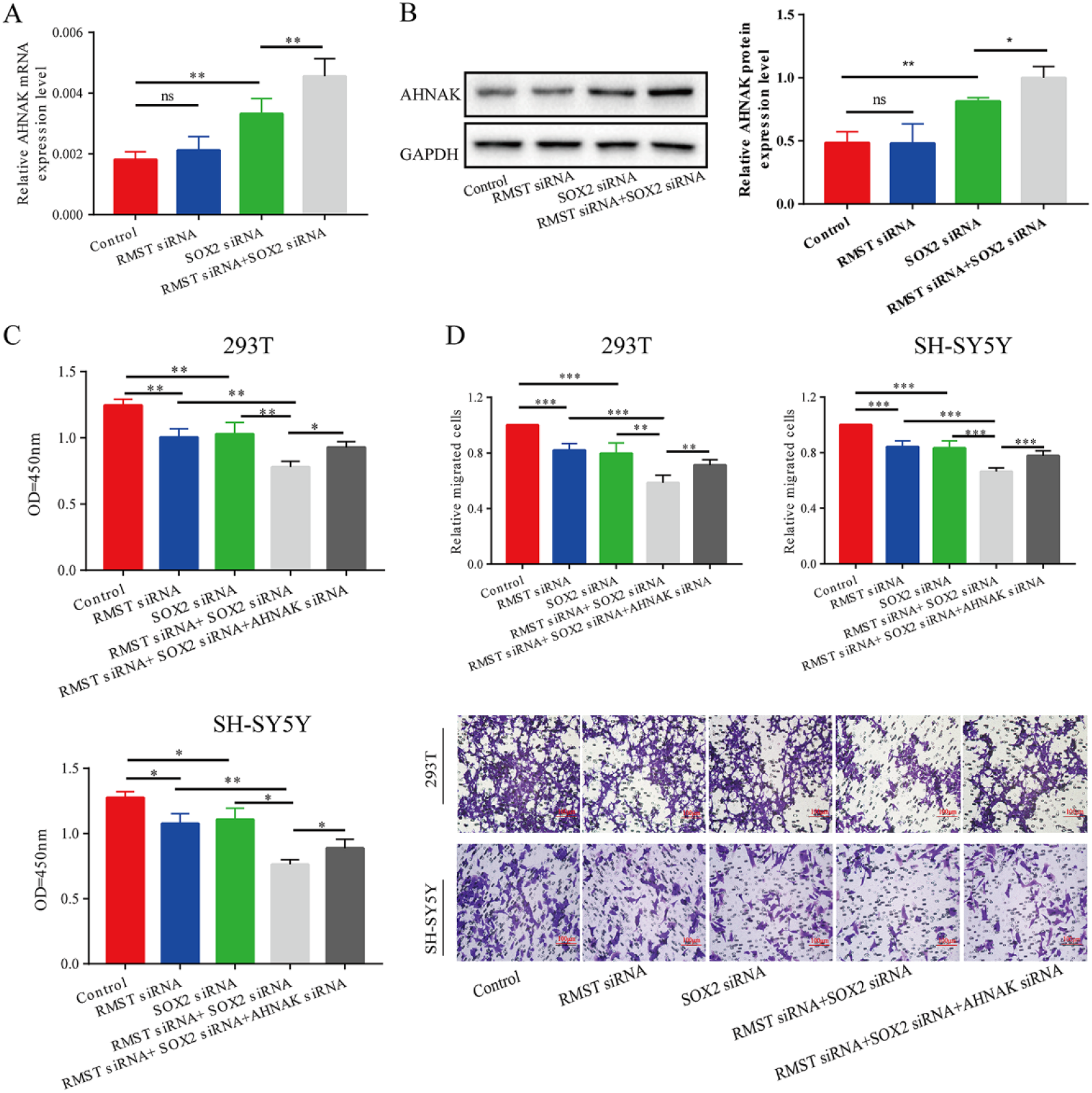
RMST played as a SOX2 transcription co-regulator to inhibit miR-1251 and raise AHNAK expression. **A and B** The mRNA and protein level of AHNAK in every group. **C and D** CCK-8 and Transwell assays showed that the combined inhibition of both RMST and SOX2 low expression on cell proliferation and migration could be partially alleviated by simultaneously downregulating the expression of AHNAK. ns *P*<0.05, \**P*<0.05, \*\**P*<0.01, \*\*\**P*<0.001

## 4. DISCUSSION

Non coding RNA (ncRNA) has been once considered as the “noise” of human genome transcription, which has no biological effects. With the development of high-throughput sequencing technology and in-depth research, more and more ncRNAs, especially miRNA and lncRNA, have been found to play very important roles in epigenetic regulation and take part in the occurrence and development of multiple diseases(Moradimotlagh et al., 2019; Pan et al., 2020; Tai et al., 2020). Although there have been some reports about ncRNAs in HSCR, its mode of action and mechanism still need further study(Gunadi et al., 2019; Zhi et al., 2018). In this study, our research team detected that in the stenosis tract of HSCR patients, RMST and miR-1251 were down-regulated apparently. MiR-1251 was firstly demonstrated as a potential prognostic markers in head and neck squamous cell carcinoma(Hui et al., 2016). But there’re few studies about miR-1251 in other diseases. Herein, it was initially found that the cell proliferation and migration was significantly inhibited after the expression of RMST or miR-1251 was reduced, indicating that RMST and miR-1251 might play a certain role in the pathogenesis of HSCR. Although miR-1251 derived from the intron region of RMST, we found RMST did not regulate the expression of miR-1251 independently. Whether there’s other regulatory mechanism? We explored it in the next step.

According to the bioinformatical analysis, we found SOX2, a transcription factor, probably bind to the promoter region of miR-1251. ChIP assay then confirmed it. SOX family, such as SOX10 has been proved to be related to the pathogenesis of HSCR(Southard-Smith et al., 1998), but there are few reports about the role and mechanism of SOX2 in the occurrence of HSCR. As reported before, SOX2 could regulate the proliferation and differentiation of peripheral nerve cells in the peripheral nervous system, (Wakamatsu et al., 2004). A recent study revealed that SOX2 gene was involved in the development of embryonic neural tube and neural crest cells(Iida et al., 2020). When knocked out SOX2 gene, the number of neurons in the ganglion derived from neural crest of mouse embryo decreased obviously, indicating SOX2 probably also play an important role in the progression of HSCR (Cimadamore et al., 2011). Furtherly, SOX2 could bind to bivalently marked promoters of poised pro-neural and neurogenic genes, and then activated neuronal differentiation appropriately (Amador-Arjona et al., 2015). These findings all indicated the important role of SOX2 played in the nervous system. In addition, over-expressed SOX2 promoted tumor progression by enhancing the abilities of cell proliferation and migration(Liu et al., 2017; Wang et al., 2017). Herein, we demonstrated SOX2 was significantly down-regulated in aganglionic tract and the following experiments showed the lowly expressed SOX2 might be involved in the occurrence of HSCR by repressing neural crest cells’ proliferation and migration via regulating miR-1251.

LncRNAs could bind with some proteins to influence their function. For example, lncRNA EPIC1 accelerated the regulation of MYC on downstream genes through combing with MYC(Wang et al., 2018). In this study, RIP results showed that RMST and SOX2 had binding relationship, which provided a basis for RMST as a transcription co-regulator of SOX2 to regulate miR-1251. We also found that the inhibition on cell proliferation and migration was more obvious in knocked down both RMST and SOX2 than abated RMST or SOX2 expression alone. Furtherly, raised miR-1251 partially alleviated the combined effect of RMST and SOX2 which confirmed that RMST might play a role as a transcription co-regulatory factor of SOX2 by enhancing the regulation of SOX2 on miR-1251. Previous studies about lncRNAs in HSCR have mostly focused on the mechanism of competitive endogenous RNA (ceRNA) (Li et al., 2018; Su et al., 2018). This research, however, firstly explored the mechanism of HSCR from the perspective of lncRNA binding protein, and expanded a new direction of HSCR research.

MiRNAs generally perform their functions by degrading their target genes(Chopra et al., 2020). It was confirmed in this study that AHNAK was the target gene of miR-1251 through the analysis of biological information and the experiment of dul-luciferase reporter assay. AHNAK, as a kind of scaffold protein, is involved in the regulation of Ca2^+^ channel and the formation of actin cytoskeleton, which has a profound impact on cell migration function(Lee et al., 2014). For example, it has been found that AHNAK is significantly under-expressed in breast cancer and *in vitro* experimental results show that the proliferation and migration ability of cells is significantly impaired after the up-regulation of AHNAK(Chen et al., 2017). Owing to the expression level of AHNAK in the stenosis tract of HSCR was significantly higher than control ones, whether miR-1251 functioned through AHNAK was investigated in this study. As expected, miR-1251 inhibited cell proliferation and migration, but improved AHNAK expression could partly reverse it.

Thus, whether RMST, as a co transcription regulator of SOX2, up-regulated the expression of AHNAK by regulating miR-1251 and exerted its roles through RMST/SOX2/miR-1251/AHNKA axis needed deeply study. As results showed, none significant difference of AHNAK expression between si-RMST group and control group was found, which also confirmed that RMST did not regulate miR-1251 alone. But the expression of AHNAK improved more a lot when co-reduced RMST and SOX2 than only down-regulating SOX2. Meanwhile, the results of cell function experiments revealed that the down-regulation of AHNAK partially reversed the inhibition of RMST and SOX2 simultaneous down-regulation on cell proliferation and migration, indicating that RMST, as a co-transcription regulator of SOX2, could affect the expression of downstream gene AHNAK through miR-1251 and then influence neural cells’ migration and proliferation.

To sum up, this study revealed that RMST functioned as a co-transcription regulator of SOX2 to upregulate the expression of downstream gene AHNAK by strengthening the regulation of SOX2 on miR-1251 for the first time, which probably be involved in the pathogenesis of HSCR. The discovery of RMST/SOX2/miR-1251/AHNAK could be helpful for the targeted therapy of HSCR in the future.

However, this study. still existed some deficiencies We discovered that decreased the expression of RMST alone could also inhibit cell proliferation and migration, whether RMST has other regulation pattern needs further study. In addition, due to the animal model of HSCR is hard to be established, this study is not supported by *in vivo* experiments, though we’re trying to overcome this shortage to make our study more persuasive.

## ACKNOWLEDGMENTS

We thank Dr. Jie Zhang, Xiaofeng Lv, Weiwei Jiang, Huan Chen, Wei Li, Changgui Lu (Children’s Hospital of Nanjing Medical University) for sample collection. This work was supported by the Natural Science Foundation of China (NSFC 81701493).

## ETHICS APPROVAL AND CONSENT TO PARTICIPATE

This study was approved by the Institutional Ethics Committee of Nanjing Medical University (NJMU Birth Cohort), and the experiments were conducted in accordance with the principles of the Declaration of Helsinki. All parents of patients had provided written informed consent in the study.

---

**Fig. S1.**
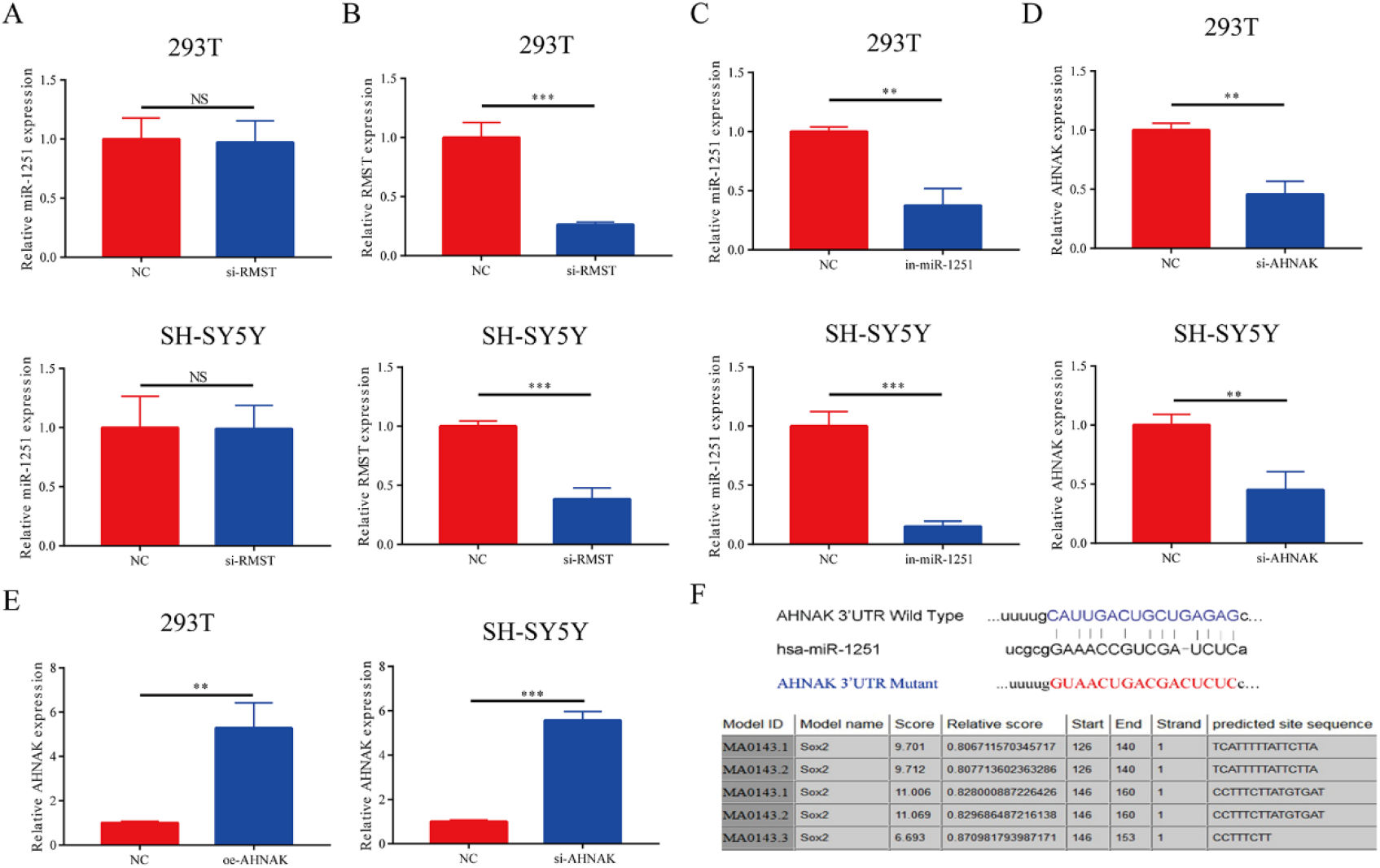
**A** The down-regulation of RMST has no significant effect on the expression level of miR-1251 in 293T and SY5Y cells. The transfection efficiency of si-RMST **B** miR-1251 inhibitor **C** si-AHNAK **D** and oe-AHNAK **E** in 293T and SY5Y cells. **F** Bioinformatic analysis predicted the potential target sites between miR-1251 and AHNAK, and SOX2 was predicted to bind with the 2kbp upstream promoter region of miR-1251. ns *P*<0.05, \*\**P*<0.01, \*\*\**P*<0.001

## REFERENCES

Alqadah, A., Hsieh, Y. W., Vidal, B., Chang, C., Hobert, O. and Chuang, C. F. (2015). Postmitotic diversification of olfactory neuron types is mediated by differential activities of the HMG-box transcription factor SOX-2. EMBO J 34, 2574–89.

Amador-Arjona, A., Cimadamore, F., Huang, C. T., Wright, R., Lewis, S., Gage, F. H. and Terskikh, A. V. (2015). SOX2 primes the epigenetic landscape in neural precursors enabling proper gene activation during hippocampal neurogenesis. Proc Natl Acad Sci U S A 112, E1936–45.

Bergeron, K. F., Silversides, D. W. and Pilon, N. (2013). The developmental genetics of Hirschsprung’s disease. Clin Genet 83, 15–22.

Cao, F., Wang, Z., Feng, Y., Zhu, H., Yang, M., Zhang, S. and Wang, X. (2020). lncRNA TPTEP1 competitively sponges miR3285p to inhibit the proliferation of nonsmall cell lung cancer cells. Oncol Rep.

Chen, B., Wang, J., Dai, D., Zhou, Q., Guo, X., Tian, Z., Huang, X., Yang, L., Tang, H. and Xie, X. (2017). AHNAK suppresses tumour proliferation and invasion by targeting multiple pathways in triple-negative breast cancer. J Exp Clin Cancer Res 36, 65.

Cheng, H., Sun, M., Wang, Z. L., Wu, Q., Yao, J., Ren, G. and Sun, X. L. (2020). LncRNA RMST-mediated miR-107 transcription promotes OGD-induced neuronal apoptosis via interacting with hnRNPK. Neurochem Int 133, 104644.

Chopra, N., Wang, R., Maloney, B., Nho, K., Beck, J. S., Pourshafie, N., Niculescu, A., Saykin, A. J., Rinaldi, C., Counts, S. E. et al. (2020). MicroRNA-298 reduces levels of human amyloid-beta precursor protein (APP), beta-site APP-converting enzyme 1 (BACE1) and specific tau protein moieties. Mol Psychiatry.

Cimadamore, F., Fishwick, K., Giusto, E., Gnedeva, K., Cattarossi, G., Miller, A., Pluchino, S., Brill, L. M., Bronner-Fraser, M. and Terskikh, A. V. (2011). Human ESC-derived neural crest model reveals a key role for SOX2 in sensory neurogenesis. Cell Stem Cell 8, 538–51.

Collignon, J., Sockanathan, S., Hacker, A., Cohen-Tannoudji, M., Norris, D., Rastan, S., Stevanovic, M., Goodfellow, P. N. and Lovell-Badge, R. (1996). A comparison of the properties of Sox-3 with Sry and two related genes, Sox-1 and Sox-2. Development 122, 509–20.

Gunadi, Budi, N. Y. P., Kalim, A. S., Santiko, W., Musthofa, F. D., Iskandar, K. and Makhmudi, A. (2019). Aberrant expressions of miRNA-206 target, FN1, in multifactorial Hirschsprung disease. Orphanet J Rare Dis 14, 5.

Hui, L., Wu, H., Yang, N., Guo, X. and Jang, X. (2016). Identification of prognostic microRNA candidates for head and neck squamous cell carcinoma. Oncol Rep 35, 3321–30.

Iida, H., Furukawa, Y., Teramoto, M., Suzuki, H., Takemoto, T., Uchikawa, M. and Kondoh, H. (2020). Sox2 gene regulation via the D1 enhancer in embryonic neural tube and neural crest by the combined action of SOX2 and ZIC2. Genes Cells 25, 242–256.

Jaroy, E. G., Acosta-Jimenez, L., Hotta, R., Goldstein, A. M., Emblem, R., Klungland, A. and Ougland, R. (2019). “Too much guts and not enough brains”: (epi)genetic mechanisms and future therapies of Hirschsprung disease -a review. Clin Epigenetics 11, 135.

Lee, I. H., Sohn, M., Lim, H. J., Yoon, S., Oh, H., Shin, S., Shin, J. H., Oh, S. H., Kim, J., Lee, D. K. et al. (2014). Ahnak functions as a tumor suppressor via modulation of TGFbeta/Smad signaling pathway. Oncogene 33, 4675–84.

Li, Y., Zhou, L., Lu, C., Shen, Q., Su, Y., Zhi, Z., Wu, F., Zhang, H., Wen, Z., Chen, G. et al. (2018). Long non-coding RNA FAL1 functions as a ceRNA to antagonize the effect of miR-637 on the down-regulation of AKT1 in Hirschsprung’s disease. Cell Prolif 51, e12489.

Liu, K., Xie, F., Gao, A., Zhang, R., Zhang, L., Xiao, Z., Hu, Q., Huang, W., Huang, Q., Lin, B. et al. (2017). SOX2 regulates multiple malignant processes of breast cancer development through the SOX2/miR-181a-5p, miR-30e-5p/TUSC3 axis. Mol Cancer 16, 62.

McKeown, S. J., Stamp, L., Hao, M. M. and Young, H. M. (2013). Hirschsprung disease: a developmental disorder of the enteric nervous system. Wiley Interdiscip Rev Dev Biol 2, 113–29.

Moradimotlagh, A., Arefian, E., Rezazadeh Valojerdi, R., Ghaemi, S., Jamshidi Adegani, F. and Soleimani, M. (2019). MicroRNA-129 Inhibits Glioma Cell Growth by Targeting CDK4, CDK6, and MDM2. Mol Ther Nucleic Acids 19, 759–764.

Ng, S. Y., Bogu, G. K., Soh, B. S. and Stanton, L. W. (2013). The long noncoding RNA RMST interacts with SOX2 to regulate neurogenesis. Mol Cell 51, 349–59.

Pan, J., Fang, S., Tian, H., Zhou, C., Zhao, X., Tian, H., He, J., Shen, W., Meng, X., Jin, X. et al. (2020). lncRNA JPX/miR-33a-5p/Twist1 axis regulates tumorigenesis and metastasis of lung cancer by activating Wnt/beta-catenin signaling. Mol Cancer 19, 9.

Sannino, G., Marchetto, A., Ranft, A., Jabar, S., Zacherl, C., Alba-Rubio, R., Stein, S., Wehweck, F. S., Kiran, M. M., Holting, T. L. B. et al. (2019). Gene expression and immunohistochemical analyses identify SOX2 as major risk factor for overall survival and relapse in Ewing sarcoma patients. EBioMedicine 47, 156–162.

Schaefer, T. and Lengerke, C. (2020). SOX2 protein biochemistry in stemness, reprogramming, and cancer: the PI3K/AKT/SOX2 axis and beyond. Oncogene 39, 278–292.

Schepers, G. E., Teasdale, R. D. and Koopman, P. (2002). Twenty pairs of sox: extent, homology, and nomenclature of the mouse and human sox transcription factor gene families. Dev Cell 3, 167–70.

Sergi, C. (2015). Hirschsprung’s disease: Historical notes and pathological diagnosis on the occasion of the 100(th) anniversary of Dr. Harald Hirschsprung’s death. World J Clin Pediatr 4, 120–5.

Shen, S., Wang, J., Zheng, B., Tao, Y., Li, M., Wang, Y., Ni, X., Suo, T., Liu, H., Liu, H. et al. (2019). LINC01714 Enhances Gemcitabine Sensitivity by Modulating FOXO3 Phosphorylation in Cholangiocarcinoma. Mol Ther Nucleic Acids 19, 446–457.

Southard-Smith, E. M., Kos, L. and Pavan, W. J. (1998). Sox10 mutation disrupts neural crest development in Dom Hirschsprung mouse model. Nat Genet 18, 60–4.

Su, Y., Wen, Z., Shen, Q., Zhang, H., Peng, L., Chen, G., Zhu, Z., Du, C., Xie, H., Li, H. et al. (2018). Long non-coding RNA LOC100507600 functions as a competitive endogenous RNA to regulate BMI1 expression by sponging miR128-1-3p in Hirschsprung’s disease. Cell Cycle 17, 459–467.

Sun, X. L., Wang, Z. L., Wu, Q., Jin, S. Q., Yao, J. and Cheng, H. (2019). LncRNA RMST activates TAK1-mediated NF-kappaB signaling and promotes activation of microglial cells via competitively binding with hnRNPK. IUBMB Life 71, 1785–1793.

Tai, F., Gong, K., Song, K., He, Y. and Shi, J. (2020). Enhanced JunD/RSK3 signalling due to loss of BRD4/FOXD3/miR-548d-3p axis determines BET inhibition resistance. Nat Commun 11, 258.

Tam, P. K. (2016). Hirschsprung’s disease: A bridge for science and surgery. J Pediatr Surg 51, 18–22.

Tang, W., Tang, J., He, J., Zhou, Z., Qin, Y., Qin, J., Li, B., Xu, X., Geng, Q., Jiang, W. et al. (2015). SLIT2/ROBO1-miR-218-1-RET/PLAG1: a new disease pathway involved in Hirschsprung’s disease. J Cell Mol Med 19, 1197–207.

Wakamatsu, Y., Endo, Y., Osumi, N. and Weston, J. A. (2004). Multiple roles of Sox2, an HMG-box transcription factor in avian neural crest development. Dev Dyn 229, 74–86.

Wang, Y., Zhou, J., Wang, Z., Wang, P. and Li, S. (2017). Upregulation of SOX2 activated LncRNA PVT1 expression promotes breast cancer cell growth and invasion. Biochem Biophys Res Commun 493, 429–436.

Wang, Z., Yang, B., Zhang, M., Guo, W., Wu, Z., Wang, Y., Jia, L., Li, S., Cancer Genome Atlas Research, N., Xie, W. et al. (2018). lncRNA Epigenetic Landscape Analysis Identifies EPIC1 as an Oncogenic lncRNA that Interacts with MYC and Promotes Cell-Cycle Progression in Cancer. Cancer Cell 33, 706–720 e9.

Wester, T. and Granstrom, A. L. (2017). Hirschsprung disease-Bowel function beyond childhood. Semin Pediatr Surg 26, 322–327.

Xu, Z., Liu, C., Zhao, Q., Lu, J., Ding, X., Luo, A., He, J., Wang, G., Li, Y., Cai, Z. et al. (2020). Long non-coding RNA CCAT2 promotes oncogenesis in triple-negative breast cancer by regulating stemness of cancer cells. Pharmacol Res 152, 104628.

Zeng, J., Li, Y., Wang, Y., Xie, G., Feng, Q., Yang, Y. and Feng, J. (2020). lncRNA 00312 Attenuates Cell Proliferation and Invasion and Promotes Apoptosis in Renal Cell Carcinoma via miR-34a-5p/ASS1 Axis. Oxid Med Cell Longev 2020, 5737289.

Zhao, J., Zhu, Y., Xie, X., Yao, Y., Zhang, J., Zhang, R., Huang, L., Cheng, J., Xia, H., He, J. et al. (2019). Pleiotropic effect of common PHOX2B variants in Hirschsprung disease and neuroblastoma. Aging (Albany NY) 11, 1252–1261.

Zhi, Z., Zhu, H., Lv, X., Lu, C., Li, Y., Wu, F., Zhou, L., Li, H. and Tang, W. (2018). IGF2-derived miR-483-3p associated with Hirschsprung’s disease by targeting FHL1. J Cell Mol Med 22, 4913–4921.

